# The *Pseudomonas aeruginosa sphBC* genes are important for growth in the presence of sphingosine by promoting sphingosine metabolism

**DOI:** 10.1101/2024.09.03.611043

**Authors:** Pauline DiGianivittorio, Lauren A. Hinkel, Jacob R. Mackinder, Kristin Schutz, Eric A. Klein, Matthew J. Wargo

## Abstract

Sphingoid bases, including sphingosine, are important components of the antimicrobial barrier at epithelial surfaces where they can cause growth inhibition and killing of susceptible bacteria. *Pseudomonas aeruginosa* is a common opportunistic pathogen that is less susceptible to sphingosine than many Gram-negative bacteria. Here, we determined that deletion of the *sphBCD* operon reduced growth in the presence of sphingosine. Using deletion mutants, complementation, and growth assays in *P. aeruginosa* PAO1, we determined that the *sphC* and *sphB* genes, encoding a periplasmic oxidase and periplasmic cytochrome c, respectively, were important for growth on sphingosine, while *sphD* was dispensable under these conditions. Deletion of *sphBCD* in *P. aeruginosa* PA14, *P. protegens* Pf-5, and *P. fluorescens* Pf01 also showed reduced growth in the presence of sphingosine. The *P. aeruginosa sphBC* genes were also important for growth in the presence of two other sphingoid bases, phytosphingosine and sphinganine. In wild-type *P. aeruginosa*, sphingosine is metabolized to an unknown non-inhibitory product, as sphingosine concentrations drop in the culture. However, in the absence of *sphBC*, sphingosine accumulates, pointing to SphC and SphB as having a role in sphingosine metabolism. Finally, metabolism of sphingosine by wild-type *P. aeruginosa* protected susceptible cells from full growth inhibition by sphingosine, pointing to a role for sphingosine metabolism as a public good. This work shows that metabolism of sphingosine by *P. aeruginosa* presents a novel pathway by which bacteria can alter host-derived sphingolipids, but it remains an open question whether SphB and SphC act directly on sphingosine.

## Introduction

In addition to their various cellular and signaling functions, some sphingolipids are key antimicrobial lipids with activity against both Gram-positive and Gram-negative bacteria(1–6). Antimicrobial sphingolipids are found at sites throughout the body including the lungs, the skin, and all mucosal surfaces(4, 7–13). Imbalances or deficiencies in barrier-associated sphingolipids, particularly sphingoid bases (examples in **Fig 1A**), increase chances of bacterial infection, illustrating the importance of these sphingolipids in defense against pathogens(14–16). The initial antibacterial action for sphingoid bases is predicted to be bacterial membrane disruption, due to their amphiphilic and detergent-like properties, followed by accumulation of sphingolipids in the cytosol, ultimately leading to cell death in both Gram-negative and Gram-positive bacteria(4, 6, 17).

**Figure 1:**
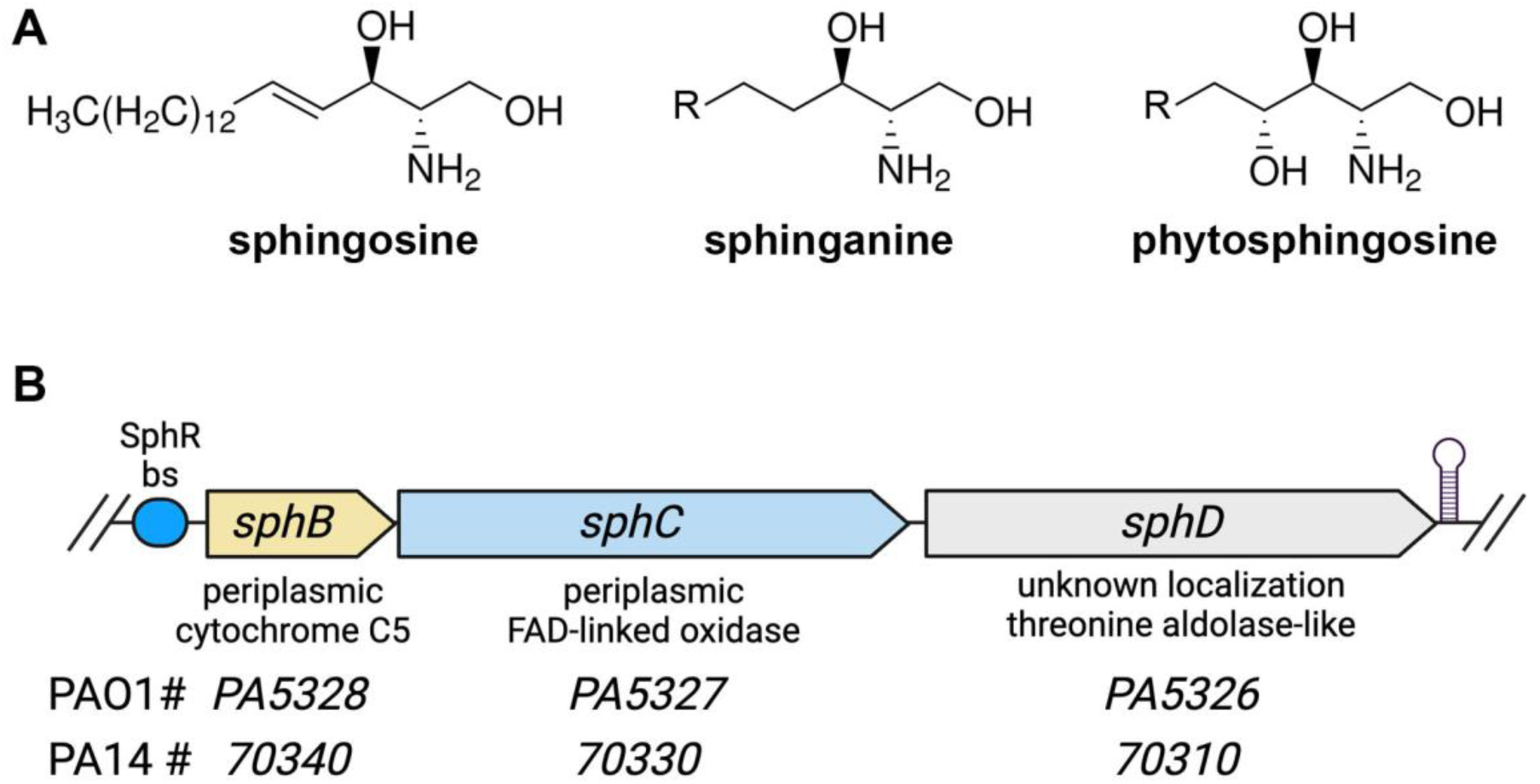
Sphingoid bases and arrangement of the *sphBCD* operon. **(A)** Structures of the sphingoid bases used in this study noting the head-group differences. All of the sphingoid bases used in this study are C18 versions, though there is tail length variation in naturally occurring versions from different body sites or organisms. **(B)** Organization of the *sphBCD* operon in *P. aeruginosa* showing the relative gene sizes, predicted functions, and the gene numbers in the PAO1 and PA14 genome. The SphR bs denotes the binding site for the sphingosine-responsive transcriptional activator SphR and the hairpin at the right edge is the predicted rho-independent terminator.

In Gram-negative bacteria, sphingolipid exposure causes separation of the inner and outer membranes, similar to the type of damage caused by cationic peptides like cathelicidins(6). The concentrations of sphingoid bases needed to cause severe and cytotoxic membrane disruption in many bacteria is low, with *Serratia marcescens* and *Pseudomonas aeruginosa* as exceptions, requiring higher concentrations or specific media conditions. For example, the minimum bactericidal concentration for *P. aeruginosa* in most media is > 1 mM, more than 300-fold higher than for *Staphylococcus aureus*, which often co-infect in lungs and wounds(1), though *P. aeruginosa* killing can be seen with concentrations as low as 10 µM under distinct media and sphingoid base solubilization regimes(17) and sphingosine-dependent killing of *P. aeruginosa* can also be seen intracellularly(18). Although there are many factors that influence antimicrobial-bacterial interactions, the sphingolipid resistance profile of *P. aeruginosa* suggests that it possesses specific mechanisms for resistance to or detoxification of sphingoid bases.

*P. aeruginosa* is associated with a variety of infections, including hospital-acquired and ventilator-associated pneumonia and bacteremia(19–22), as well as chronic lung infection in individuals with cystic fibrosis (CF) and chronic obstructive pulmonary disorder (COPD)(22–28). Many of these infection niches contain abundant sphingosine, other sphingoid bases, and the sphingosine precursors sphingomyelin and ceramide(2, 29–33), though a decrease in sphingosine concentration due to ceramide accumulation has been shown in CF(34, 35). Therapeutic intervention to treat ceramide accumulation can rescue the susceptibility to *P. aeruginosa* infection in animal models(36). Within the context of pulmonary infections, *P. aeruginosa*’s ability to resist the antimicrobial effects of sphingosine is correlated with a survival advantage, due in part to the presence of sphingoid bases within the lung epithelium(23).

Exposure of *P. aeruginosa* to pulmonary surfactant leads to induction of a small set of sphingosine-responsive genes in an SphR-dependent manner, including a metabolic operon, *sphBCD,* encoding a predicted cytochrome c (*sphB*), predicted oxidoreductase enzyme (*sphC*), and predicted PLP-dependent aldolase enzyme (*sphD*)(23) (**Fig 1B**). We previously showed that loss of *sphC* led to a small but statistically significant reduction in *P. aeruginosa* survival in the presence of sphingosine(23). However, the conditions needed for sphingosine killing of *P. aeruginosa* are very specific. Thus, we sought to examine the effects of sphingosine conditions that may more closely mimic some infections sites. Here we demonstrate that, in addition to killing under specific conditions, sphingosine can strongly suppress growth of an *P. aeruginosa sphBCD* mutant, with follow-up experiments supporting the *sphBC* genes as important for *P. aeruginosa* growth in the presence of sphingosine via sphingosine detoxification. Sphingosine detoxification can function as a public good promoting growth of sphingosine-susceptible *P. aeruginosa* mutants.

## Results

### The importance of *sphBCD* genes for *P. aeruginosa* growth in the presence of sphingosine and sphingosine analogs

We previously reported the importance of *sphR* and *sphA* for resistance to sphingosine-dependent killing of *P. aeruginosa* PAO1 with a minor impact of *sphC* mutation(23). Killing of *P. aeruginosa* by sphingosine requires specific media conditions (high divalent cation concentration) and/or micellular sphingosine(17, 23). We observed that even in the absence of these very particular conditions and thus the absence of killing, sphingosine could strongly inhibit growth of *P. aeruginosa* Δ*sphBCD* deletion mutants and that inhibition was stronger when growth was conducted in glass rather than in plastic at a given concentration of sphingosine (**Fig 2A**). The same protective role of *sphBCD* can be observed during growth in the presence of sphinganine (**Fig 2B**) and phytosphingosine (**Fig 2C**).

**Figure 2:**
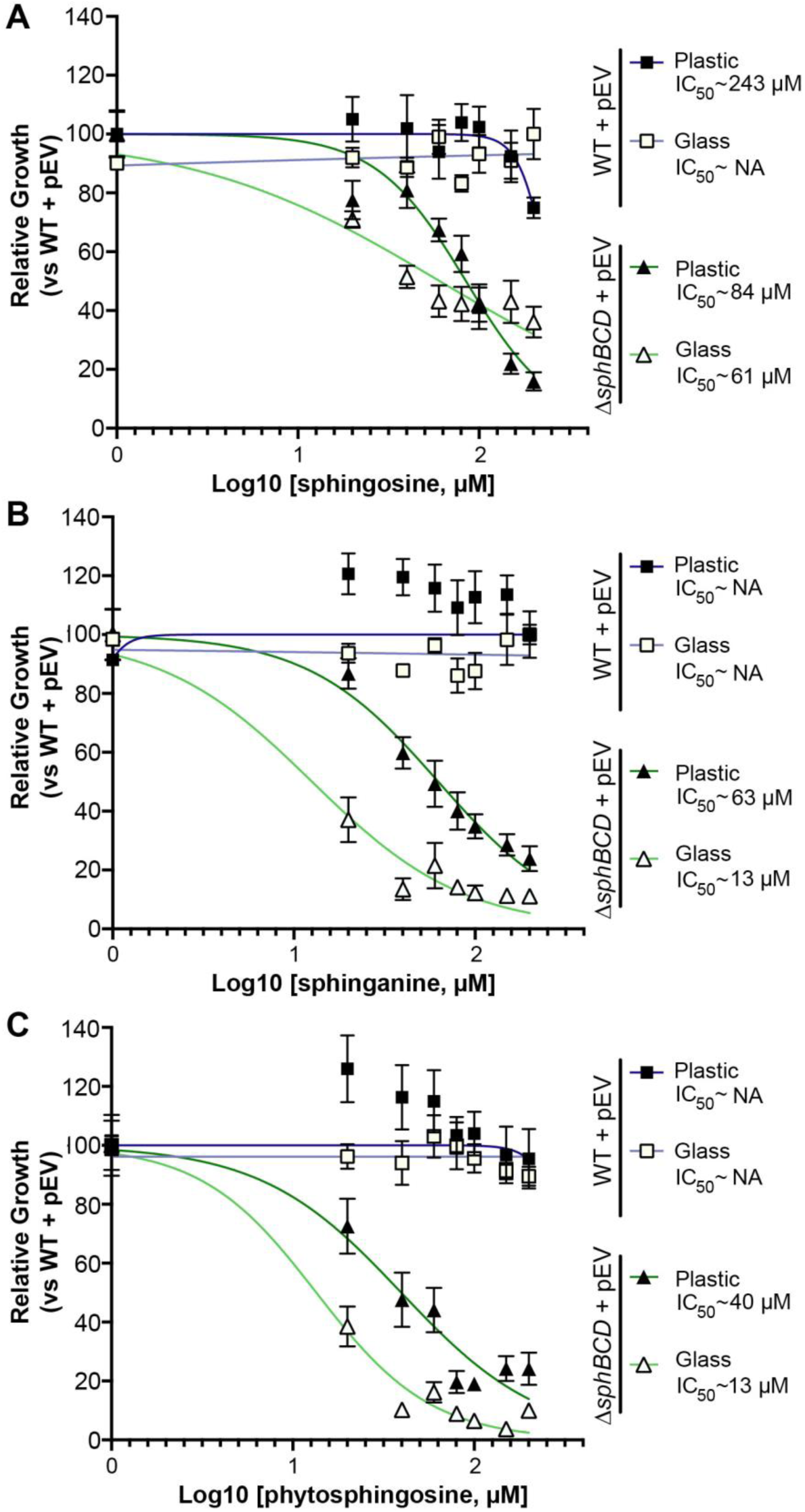
Concentration dependent inhibition for sphingoid bases is dependent on the base and the culture vessel material. All panels show relative growth measured by OD_600_ as compared to the WT with empty vector (pEV) in the absence of sphingosine at the 18-hour timepoint. The data shown here are for **(A)** sphingosine, **(B)** sphinganine, and **(C)** phytosphingosine in either glass (open symbols) or plastic (closed symbols). The IC_50_ curve and estimated IC_50_s to the right of the plots generated using variable-slope curve fitting in GraphPad Prism. If calculated IC_50_ was above the solubility of sphingosine, it was listed as NA. Data points denote means summarizing three independent experiments and error bars mark standard deviation. Abbreviations: pEV, empty vector pMQ80.

Deletion of the *sphBCD* operon increased susceptibility to sphingosine and close analogs when measured at 18 hours post inoculation (**Fig 2**), and we wanted to examine the kinetics of this growth inhibition by measuring growth over time. We measured growth with 100% set as WT OD600 in the absence of sphingosine at 18 hours. At 200 µM sphingosine, the *sphBCD* deletion shows initial growth that starts to plateau after about 10 hours, while WT has a delay in growth before resumption of a nearly normal growth rate. The complementation strain has no substantial delay, growing at a rapid rate after lag phase (**Fig 3A**). The *sphBCD* deletion is also defective for growth in the presence of sphinganine (**Fig 3B**) and phytosphingosine (**Fig 3C**). Neither of these sphingosine analogs shows the strong delay in WT growth and, while Δ*sphBCD* growth in phytosphingosine shows the same plateau as for sphingosine (compare **Fig 3C** with **3A**), the Δ*sphBCD* strain can grow slowly in the presence of sphinganine with a substantial delay. Growth of all strains in the absence of sphingosine is presented in **Supplemental Figure S1**.

**Figure 3:**
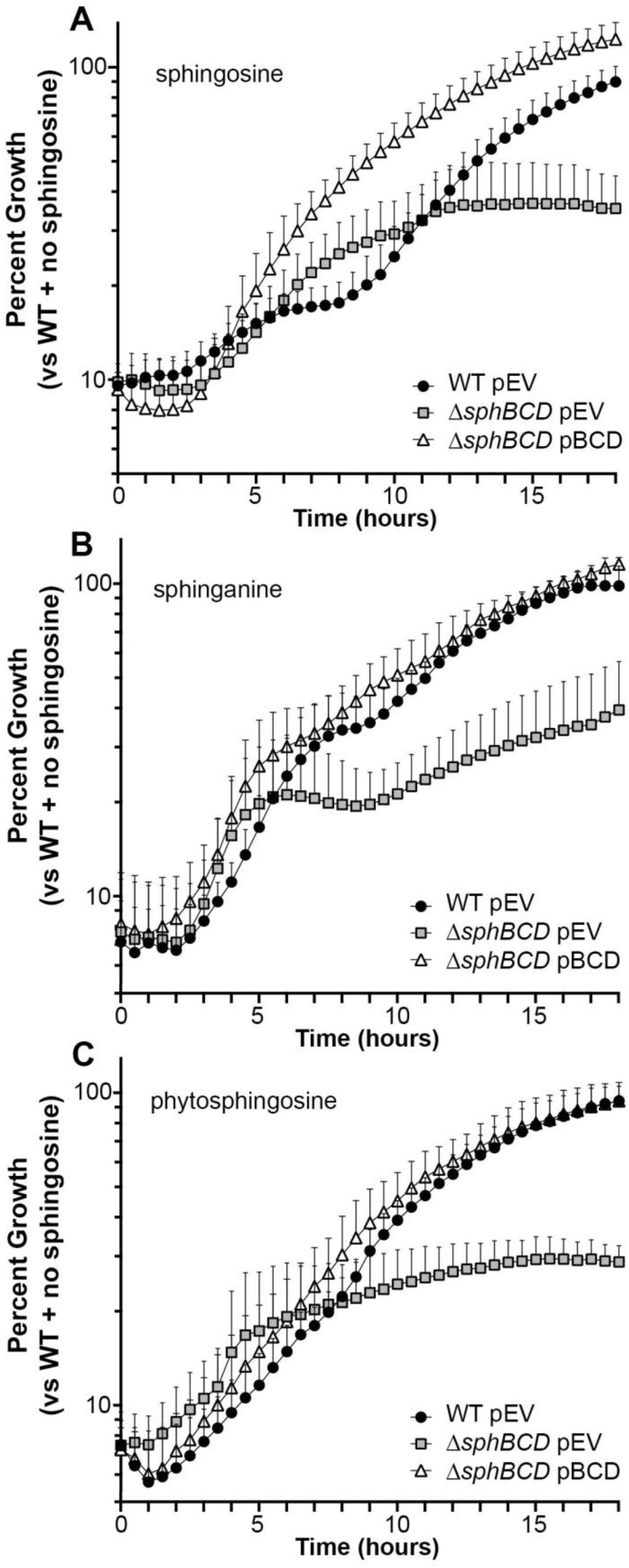
Kinetic growth assessment of wild-type, mutant, and complemented strains in the presence of sphingoid bases. All panels are 18-hour timecourses measuring relative growth of each strain at each timepoint as measured by OD_600_ compared to the WT with empty vector (pEV) in the absence of sphingosine at the 18h timepoint. The data shown here are for **(A)** sphingosine, **(B)** sphinganine, and **(C)** phytosphingosine. Growth curves in MOPS pyruvate in the absence of sphingoid bases are presented in Supplemental Figure S1. Data points denote means summarizing three independent experiments and error bars mark standard deviation with only the bars above the mean shown for figure clarity. Abbreviations: pEV, empty vector pMQ80; pBCD, vector containing *sphBCD*.

### The critical role for *sphC* for growth in the presence of sphingosine

Deletion of *sphBCD* can be complemented by plasmids carrying *sphBCD* or a plasmid carrying *sphBC*, but not other single genes from the locus (**Fig 4A**), supporting *sphB* and *sphC* as required components for resistance to sphingosine. Deletion of *sphC* alone phenocopies Δ*sphBCD* and *sphC* complements this phenotype in Δ*sphC* (**Fig 4B**). Similar to sphingosine, deletion of *sphC* results in growth inhibition by the sphingosine analogs sphinganine and phytosphingosine, and these phenotypes can be complemented (**Fig 4C & D**). These data support a role for *sphBC* in resistance to growth inhibition by antimicrobial sphingoid bases.

**Fig 4.**
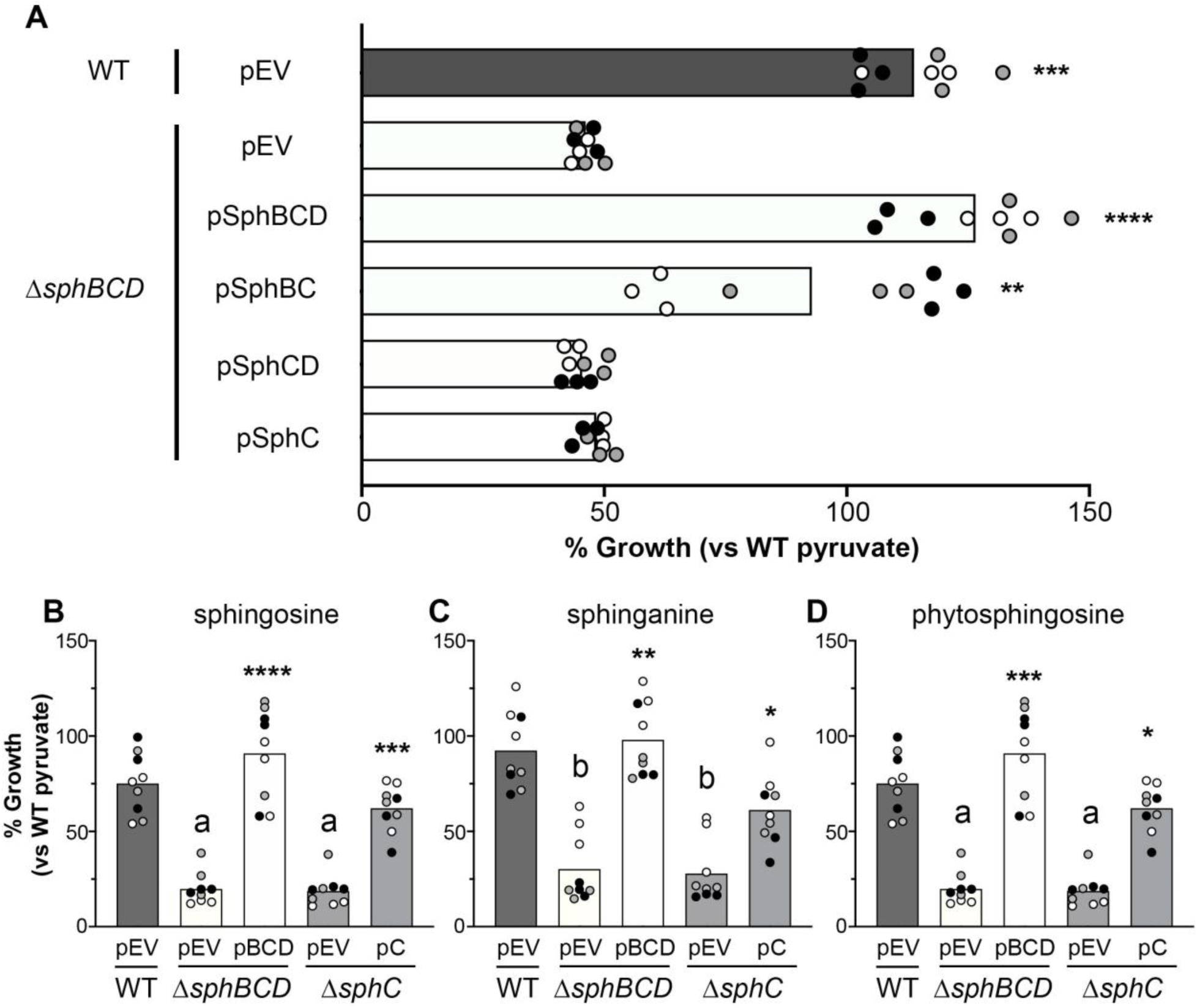
The *sphBC* genes are critical for wild-type levels of growth in the presence of sphingoid bases. **(A)** Complementation analysis of Δ*sphBCD* shows significant complementation only with plasmids expressing *sphB* and *sphC*, while *sphD* appears dispensable for growth at 18 hour normalized to WT empty vector growth in MOPS pyruvate set as 100%. **(B-D)** Deletion of *sphC* phenocopies deletion of *sphBCD* and can be complemented by *sphC* on a plasmid. This phenotype is shared between the sphingoid bases **(B)** sphingosine, **(C)** sphinganine, and **(D)** phytosphingosine. 18-hour growth was normalized to WT empty vector growth in MOPS pyruvate set as 100%. For all panels, all data points are shown and are colored by experiment with white circles for all replicates from experiment #1, gray from experiment #2, and black from experiment #3. Only the means for each experiment are used in the statistical analyses for these panels (n = 3 per condition). For (A), significance noted as (**, p<0.01; ***, p<0.001; ****, p<0.0001) calculated from ANOVA with Dunnett’s post-test with Δ*sphBCD* pEV as the comparator. For (B-D), significance noted as (**, p<0.01; ***, p<0.001; ****, p<0.0001) for comparisons of each complemented strain to its empty vector control, while significant difference to WT with empty vector noted as (a, p<0.0001; b, p<0.01). Analysis of B-D conducted with ANOVA and Tukey’s post-test comparing all groups. Abbreviations: pEV, empty vector pMQ80; pBCD, vector containing *sphBCD;* pC, vector containing *sphC*.

### *sphBC* are important for metabolism of sphingosine to a non-toxic metabolite

While there were many potential mechanisms by which *sphBC* could provide sphingosine resistance, one potential mechanism was metabolism of sphingoid bases to a compound that was not growth inhibitory. The *sphC* gene encodes a TAT-secreted periplasmic oxidoreductase(37) and *sphB* encodes a sec-secreted periplasmic cytochrome c5-like protein, predicted to be a lipoprotein. These predicted functions suggested a role for oxidation of some compound in the periplasm, potentially sphingosine or a compound required for subsequent sphingosine metabolism. Sphingosine is depleted from supernatants and cell culture extracts of WT cells (**Fig 5** and also seen in(38)), while sphingosine and close analogs accumulate in cell culture extracts of Δ*sphBCD*, as measured by bioassay (**Fig 5A-C**). The bioassay measures are supported by liquid-chromatography mass spectrometry (**Fig 5D**) and thin-layer chromatography (TLC) (**Fig 5E**). These data support a role for *sphBC* in sphingosine metabolism to a non-toxic product, as functional *sphBC* (WT) leads to no substantial growth inhibition and absence of the added sphingosine.

**Figure 5:**
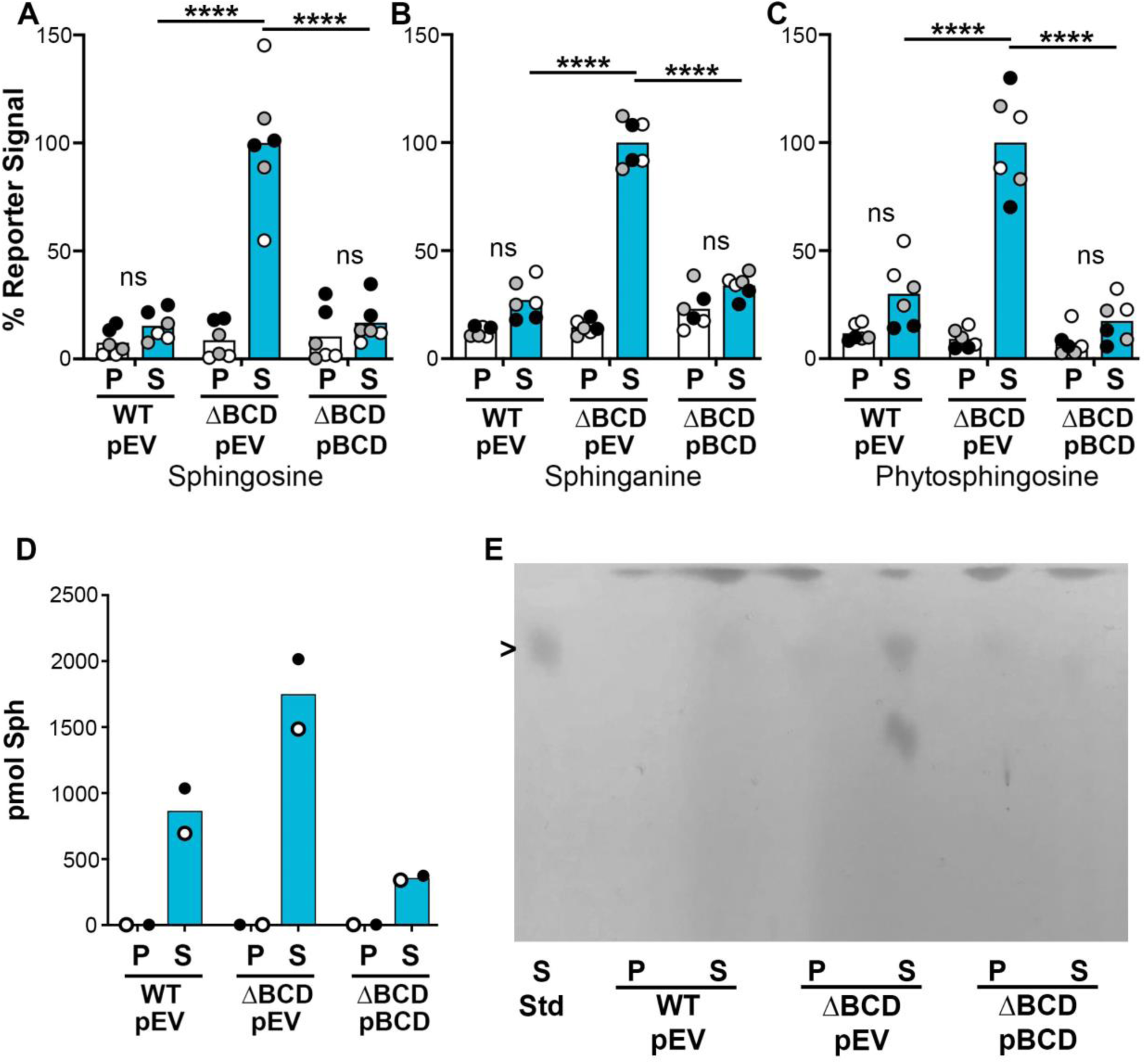
Metabolism of sphingoid bases is dependent on the presence of *sphBCD*. **(A-C)** Determination of sphingoid bases remaining in the culture after 18 hours of incubation as measured using the *sphA-lacZ* reporter assay, **(D)** LC-MS, and **(E)** TLC. Statistical significance noted as ****P < 0.0001 using a two-way ANOVA with Tukey’s post-test comparing all groups. For panels A-D, all data points are shown and are colored by experiment with white circles for all replicates from experiment #1, gray from experiment #2, and black from experiment #3. Only the means for each experiment are used in the statistical analyses for these panels (n = 3 per condition, except the LC-MS, for which only two replicates were run and are therefore not statistically analyzed). The spot that runs below sphingosine in the ΔsphBCD mutant TLC lane (in E) is an unknown amine-containing lipid and did not run similarly to any of our sphingolipid standards. Abbreviations: ns, not significant; P, pyruvate (control); S, sphingosine; ΔBCD, Δ*sphBCD*; pEV, empty vector pMQ80; pBCD, vector containing *sphBCD*.

### Phylogenetic distribution of the *sphBCD* genes and their roles in other species

The *sphBCD* genes are present in all sequenced *P. aeruginosa* and are also present in most non-*aeruginosa* Pseudomonads using the Pseudomonas genome browser(39). As *sphB* and *sphC* encode proteins in large families, true orthology is difficult to assess, particularly in the absence of any direct understanding of substrate interaction in the case of SphC. Co-occurrence searches with String(40) yield quite a large number of hits in the Firmicutes, Actinobacteria, and Alpha-, Beta-, and Gamma-Proteobacteria, but nothing outside of those groups. Manual searching through the resultant genes suggests some could be orthologs, including a putative SphC of *Caulobacter crescentus*, described below, while others are likely unrelated to sphingosine. Therefore, we first focused on assessing function of the *sphBCD* genes in other Pseudomonads, including *P. fluorescens* WCS365 which does not have an *sphD* ortholog in its *sphBC* operon. Deletion of the *sphBCD* genes from *P. aeruginosa* PA14 and *P. protogens* Pf-5 showed a growth defect in the presence of 200 µM sphingosine regardless of culture vessel material (**Fig 6A&B**). The *sphBCD* deletion mutant of *P. fluorescens* Pf01 was lower than WT in each vessel material, but the difference was only significant in glass (**Fig 6A**). Deletion of *sphBC* in *P. fluorescens* WCS365 did not show a phenotype. These data support a role for *sphBC* in resistance to sphingosine beyond *P. aeruginosa*, but presence of these genes does not necessarily predict importance for growth in the presence of sphingosine. We deleted the *sphC* gene from *C. crescentus* but the measured effect was significant only within a very small sphingosine concentration range (**Supplemental Figure S2A**). We also tested heterologous expression of *C. crescentus sphBC* to attempt complementation of *P. aeruginosa ΔsphBCD*. For *C. crescentus* putative *sphBC*, the native sec- and TAT-signal sequences encoded in *sphB* and *sphC*, respectively, were swapped for the sec- and TAT-signal sequences from *P. aeruginosa sphB* and *sphC*. While data show a trend towards partial rescue, the impact of *C. crescentus sphBC* in *P. aeruginosa* is not statistically significant (**Supplemental Figure S2B**).

**Fig 6.**
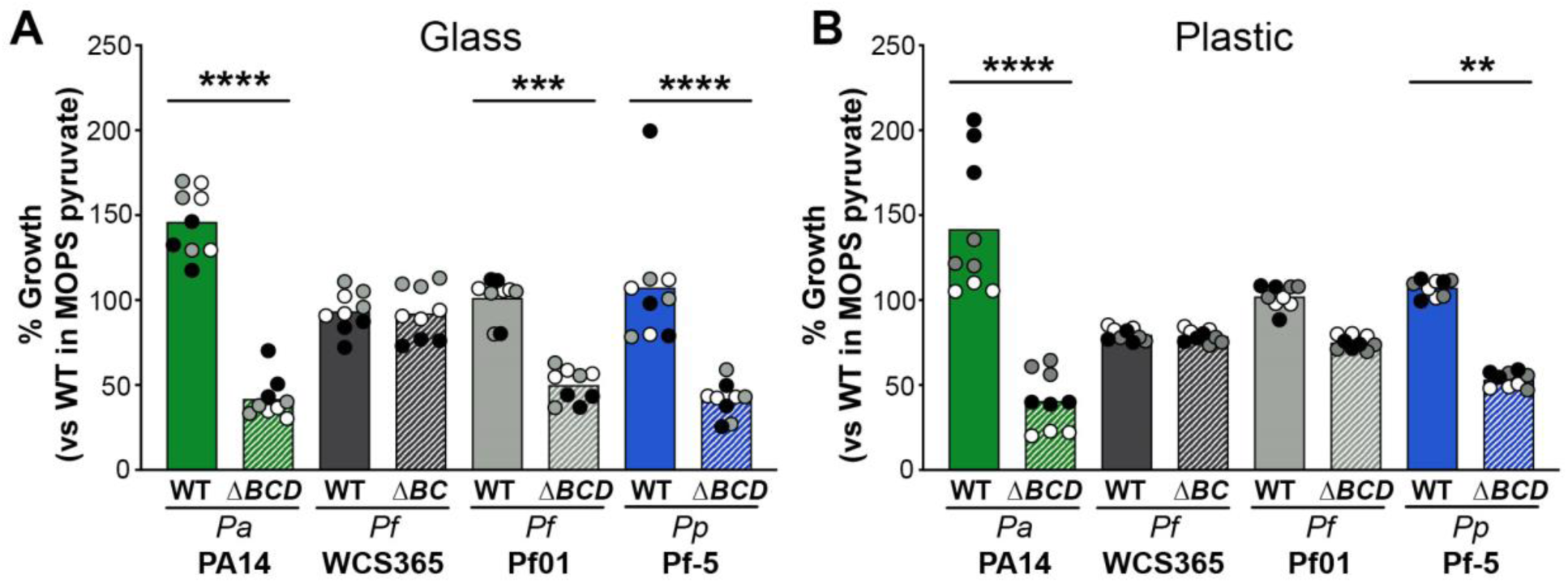
The role of *sphBC* in other Pseudomonads. The growth WT and mutant for each strain in 200 µM sphingosine shown normalized to that strain’s growth in MOPS pyruvate media. As seen for *P. aeruginosa* PAO1 (Fig 2), the culture vessel material impacts the potency of sphingosine for some strains. Significance noted as (**, p<0.01; ***, p<0.001; ****, p<0.0001) calculated from ANOVA with Sidak’s post-test comparing WT to mutant within each strain. For both panels, all data points are shown and are colored by experiment with white circles for all replicates from experiment #1, gray from experiment #2, and black from experiment #3. Only the means for each experiment are used in the statistical analyses for these panels (n = 3 per condition) Abbreviations: ΔBCD, Δ*sphBCD*; ΔBC, Δ*sphBC*; Pa, *P. aeruginosa*; Pf, *P. fluorescens*; Pp, *P. protogens*.

### Detoxification of sphingosine is a public good

The *sphBC* genes have a role in metabolism of sphingosine to a product that is not growth inhibitory, which suggests that cells capable of sphingosine metabolism could potentially protect cells that cannot otherwise metabolize sphingosine from its antimicrobial effects. Wild-type *P. aeruginosa* partially protects Δ*sphBCD* from sphingosine growth inhibition as measured by both fluorescent signal (**Fig 7A**) and CFU (**Fig 7B**) and the same effect was seen when the fluorescent markers were swapped between the strains (**Supplemental Fig S3**). *P. aeruginosa* could likewise protect the sphingosine-susceptible *Staphylococcus aureus* (**Fig 8A**). While protection of *S. aureus* by Δ*sphBCD* trended lower than wild type (**Fig 8B**), this did not reach significance given the assay variability.

**Fig 7.**
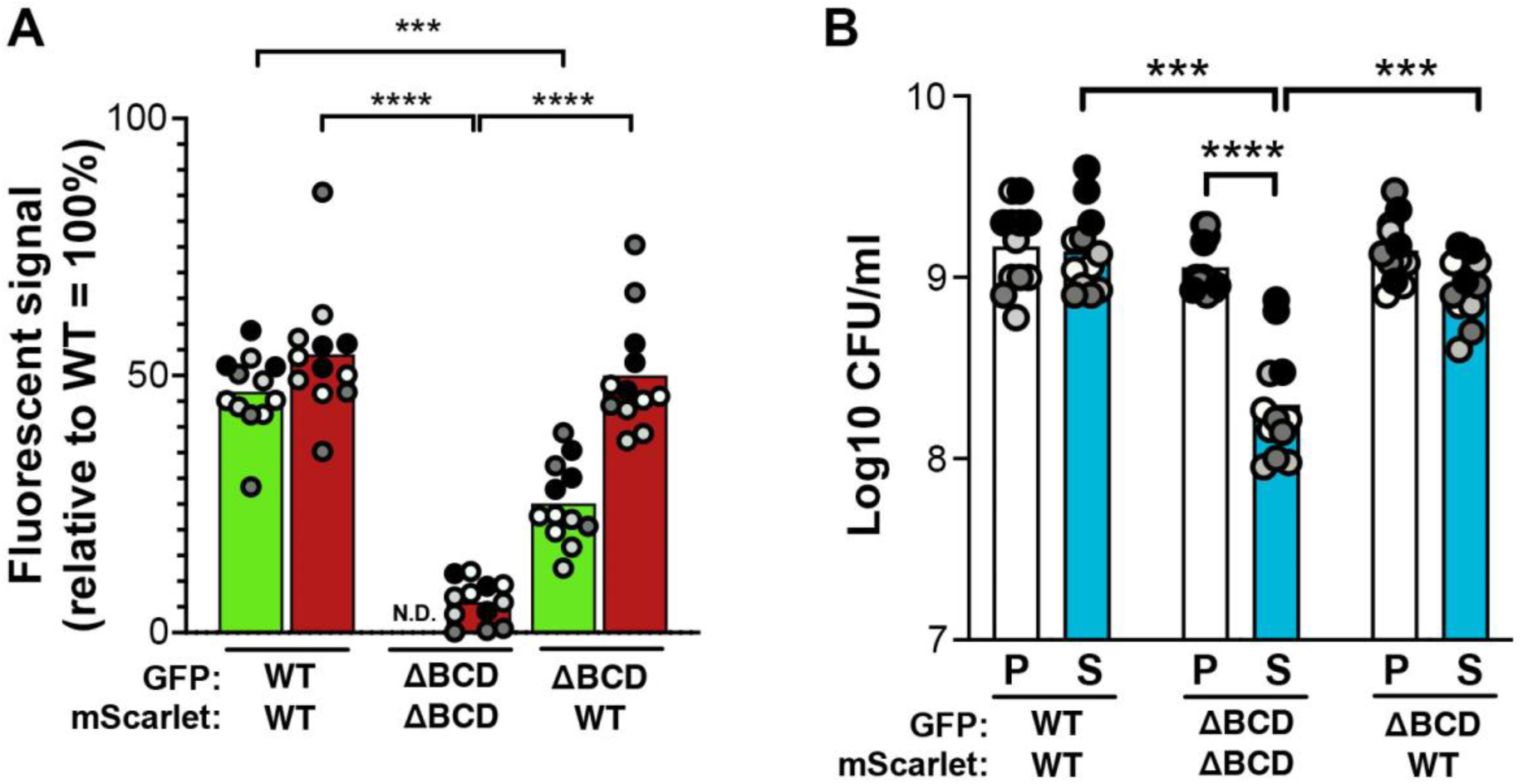
Wild type sphingosine detoxification can protect co-cultured Δ*sphBCD* from growth inhibition by sphingosine. **(A)** Fluorescence signal for GFP and mScarlet normalized to monoculture of WT carrying GFP or mScarlet, respectively. GFP signal shown with light green bars and mScarlet signal shown with dark red bars. The strain carrying each fluorescent protein is labeled below graph. **(B)** CFU counts of GFP-expressing colonies in the presence (S) or absence (P) of sphingosine. The strain carrying each fluorescent protein is labeled below graph. Significance noted as (***, p<0.001; ****, p<0.0001) calculated from ANOVA with Tukey’s post-test comparing within and between co-culture groups. For both panels, all data points are shown and are colored by experiment with white circles for all replicates from experiment #1, light gray from experiment #2, dark gray from experiment #3, and black from experiment #4. Only the means for each experiment are used in the statistical analyses for these panels (n = 4 per condition) Abbreviations: ΔBCD, Δ*sphBCD*; P, pyruvate (control); S, sphingosine; N.D., not detectable.

**Fig 8.**
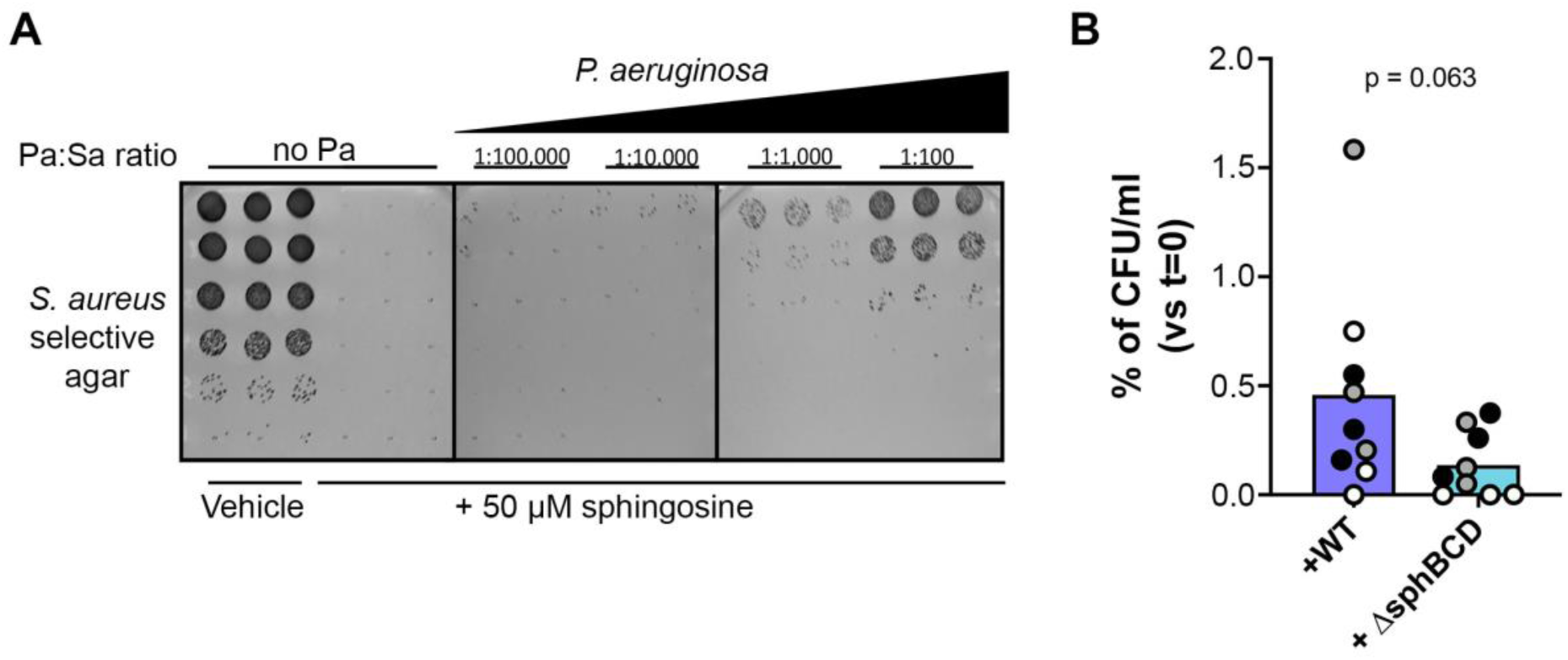
*P. aeruginosa* can protect *S. aureus* from complete killing by sphingosine. **(A)** Titration of WT *P. aeruginosa* into the sphingosine-containing media for 1h prior to *S. aureus* inoculation protected a small proportion of *S. aureus* from the lethal effects of 5 hours in the presence of 50 µM sphingosine in a *P. aeruginosa* inoculum-dependent manner. **(B)** The proportion of the initial *S. aureus* population protected by *P. aeruginosa* WT trended higher than the proportion protected by Δ*sphBCD*. All data points are shown and are colored by experiment with white circles for all replicates from experiment #1, gray from experiment #2, and black from experiment #3. Only the means for each experiment were used for a t-test to statistically analyze these data (n = 3 per condition) and thus the technical replicates with no CFU are averaged to a non-zero number for each experiment (even for experiment #1, where only one replicate had countable colonies).

## Discussion

Sphingoid bases, including sphingosine, are important antimicrobial compounds on epithelial surfaces of mammals(12, 41) and are also produced by plants and released into the rhizosphere(42). Here, we report the identification of the *P. aeruginosa sphBCD* operon as necessary for metabolism, and thus detoxification, of sphingosine and other sphingoid bases, showing that functional *sphBCD* is needed for wild-type levels of growth in the presence of sphingoid bases. These conclusions are supported by growth studies, complementation, and measurements of sphingosine metabolism. Wild-type *P. aeruginosa* can also protect susceptible bacteria from sphingosine pointing to a potential role in mixed microbial communities.

The work presented here focuses on conditions wherein sphingosine inhibits growth but is not bactericidal for either wild-type or Δ*sphBCD*. These conditions are quite different than those required for *P. aeruginosa* killing by sphingosine by us and others, which typically use very high divalent cation concentrations and is dependent on the phase of the lipid(17, 23). In our current data, sphingosine is dried onto the vessel surface allowing vehicle evaporation prior to adding media and *P. aeruginosa* and results in growth inhibition rather than killing, though others have also noted bacterial growth inhibition rather than killing for sphingoid bases(42). Therefore, while the concentration of sphingosine in the entire well is listed in our experiments, the concentration of free sphingosine in the liquid phase at any given point in time is unknown. Our model may not mimic the antimicrobial activity of sphingosine in liquid covered epithelium, like in the lung(12), and might be a closer mimic to the antimicrobial activity of sphingosine on the skin with a temporary covering of sweat(3). In a similar manner, our model is likely closer to the behavior of plant-derived sphingoid bases in non-saturated soils. The importance and properties of the vessel material underlines the difference of our model, where there are noticeable differences in concentration-dependent inhibition and growth phenotype depending on whether the culture vessel was glass or plastic. Because of the very different conditions in our model, the phenotypes shown here are not directly comparable to the killing phenotypes we previously reported(23) or to the *P. aeruginosa* killing presented by others(17). In our previous work, an *sphC* mutant had a small but measurable defect in the sphingosine killing assay while Δ*sphR* and Δ*sphA* mutants were very susceptible to sphingosine killing. However, in the growth inhibition assay, the *sphA* mutant has no phenotype (**Supplemental Figure S4A**). Additionally and interestingly, while *sphBCD* can be induced by sphingosine in an *sphR*-dependent manner(23), *sphR* is not important for growth in the presence of sphingosine (**Supplemental Figure S4B**) suggesting either that basal transcription is sufficient for growth or that there is another regulator inducing *sphBCD*, perhaps related to envelope stress response. We think that these differences in phenotypes for sphingosine-related mutants in the two sphingosine response models, killing vs growth inhibition, are likely biologically important and may reflect management of sphingosine under different environmental conditions. We also note that the carbon source in minimal media impacts the effect of sphingosine on PAO1 growth with less impact of sphingosine when grown using a carbon source which enables faster growth (**Supplemental Figure S5**), though complementation with *sphBCD* still improves growth even when there is little overall inhibition (i.e. in MOPS + Succinate). This effect of carbon source could be due to either faster growth rate or more rapid accumulation of cell mass that could dilute the effectiveness of sphingosine – these are conjecture and would need to be formally tested.

Our genetic analysis implicates SphC and SphB as the critical proteins for sphingosine resistance encoded in the *sphBCD* operon, as deletion of *sphC* phenocopies Δ*sphBCD* (**Fig 4B-D**) and only vectors containing both *sphC* and *sphB* can complement Δ*sphBCD* (**Fig 4A**). SphC is predicted to be an FMN-linked oxidoreductase that is known to be TAT secreted and localized to the periplasm(37). SphB is a predicted lipoprotein cytochrome c with a Sec signal sequence. Based on the data presented here and the presence of the *sphBCD* operon in the sphingosine:SphR regulon(23), we predict that SphC can oxidize sphingosine to a metabolite that is non-toxic and the electron needed for this oxidation is replenished by SphB. Some evidence supporting that SphC and SphB might be partners is that while plasmid-borne *sphC* can complement Δ*sphC*, it is not as strong a complementation as plasmid-borne *sphBC* complementation of Δ*sphBCD* (**Fig 4**). This could be explained by a stoichiometry mismatch between SphC and SphB. As to why the plasmid carrying *sphBC* does not complement as well as the plasmid carrying *sphBCD*, we are not sure, though since we have not measured transcript and protein levels generated from these constructs, it could simply be a difference in functional expression. It is interesting to note that the two organisms that we tested that carry only *sphBC* in an operon *P. fluorescens* WCS365 and *C. crescentus*, compared to those with *sphBCD*, show little to no effect of the *sphBC* deletion on growth in the presence of sphingosine (**Fig 6** and **Supplemental Figure S2**).

Multiple attempts to identify the direct metabolite of sphingosine were unsuccessful, perhaps because one potential initial product would be a very reactive aldehyde aldol. While our data here underscore the necessity of *sphBC* for sphingosine metabolism and normal levels of *P. aeruginosa* growth in the presence of sphingosine, we currently have no evidence that *sphBC* are sufficient for sphingosine metabolism. This leaves open the possibility that SphB and SphC act indirectly to detoxify sphingosine. Complementation of *P. aeruginosa* Δ*sphBCD* with secretion adapted *sphBC* homologs from *Caulobacter crescentus* showed no significant effect (**Supplemental Figure S2**). There are many reasons this heterologous complementation might have failed yet be non-informative, including poor protein folding in the heterologous host, secretion failure despite the attempt at secretion adaptation of each sequence to the heterologous host, rapid degradation of one or both proteins, or, in the case of SphB, failure to interact with the unknown inner membrane electron donor in the heterologous host. Additionally, while *C. crescentus* putative SphB and SphC are homologous to *P. aeruginosa* SphB (44% identity, 55% positive) and SphC (42% identity, 58% positive), it is unknown whether they are orthologous.

When we examined other strains and other Pseudomonas species, we noted that while *sphBCD* deletion led to poorer growth in the presence of sphingosine for *P. aeruginosa* PA14, *P. fluorescens* Pf01, and *P. protegens* Pf-5, deletion of *sphBC* in *P. fluorescens* WCS365 had no phenotype (**Fig 6**). Additionally, the magnitude of the phenotype differed between species and, like for *P. aeruginosa*, was dependent on the culture vessel material. Combining these findings with the observation that *P. aeruginosa* Δ*sphBCD* can still grow in the presence of sphingosine, albeit not to the same extent as wild type, we conclude that there are other proteins or cellular processes that can function independently of *sphBCD*. In *P. fluorescens* WCS365, there is no *sphD* homolog at the locus and there is very minimal decrease in growth of either wild-type or Δ*sphBC*. This strain must have an alternate mechanism to resist growth inhibition by sphingosine.

Regardless of whether SphC and SphB directly act on sphingosine, *sphBC* dependent sphingosine metabolism depletes sphingosine from the media (**Fig 5**). Such sphingosine depletion led us to hypothesize that metabolism of sphingosine by wild-type cells would protect sphingosine-susceptible bacteria in co-culture which we did indeed observe in a co-culture of wild-type and Δ*sphBCD* cells (**Fig 7**). Similarly, *S. aureus* is completely killed by 50 µM sphingosine under the conditions of our assay, but a small percentage can be protected by *P. aeruginosa*. While fewer *S. aureus* are protected by Δ*sphBCD*, variation in the means makes the contribution of *sphBCD* to this protection not statistically significant. One of the caveats of this *P. aeruginosa*-*S. aureus* co-culture is that, for these lab isolates, *P. aeruginosa* eventually kills the *S. aureus*(43–45). Future work will look at co-isolates of these species from the same patient samples, where apparently peaceful co-existence is common(46). Since many bacteria and some fungi are susceptible to sphingoid bases(3, 42), sphingoid base detoxification in areas of very high sphingosine concentration (skin, rhizoplane) might contribute to community structure and composition.

Our identification and characterization of the sphingoid base-dependent phenotype of *sphBCD* and *sphC* mutants is an important step in our understanding of bacterial manipulation of sphingolipids. However, there remain a number of important and unaddressed issues identified during our work, including the biochemical mechanism behind SphC and SphB function, the identity and role of *sphBC*-independent sphingosine management systems in *P. aeruginosa* and other Pseudomonads, and the contributions of sphingosine detoxification to spatial architecture in sessile communities.

## Materials and Methods

### Strains and growth conditions

*Pseudomonas aeruginosa* PA14, PAO1, and related mutant strains were maintained at 37°C on Pseudomonas Isolation Agar (PIA) plates with 20 μg/ml gentamicin added when appropriate. *Pseudomonas fluorescens, Pseudomonas protegens*, and strains of those species were maintained at 30 °C on lysogeny broth-Lennox formulation (LB) plates. Prior to assay set up, strains were grown shaking either at 37 °C or 30 °C overnight in a 1X MOPS medium (47), modified as previously described (48), and supplemented with 25 mM pyruvate and 5 mM glucose, adding in 20 μg/ml gentamicin when appropriate. For competition assays, *Pseudomonas aeruginosa* PAO1 and *Staphylococcus aureus* strains were maintained at 37 °C on LB plates. Prior to co-culture experiments, *P. aeruginosa* and *S. aureus* were grown shaking at 37 °C in 1X MOPS medium with 20 mM pyruvate and 5 mM glucose. All strains are listed in Table 1.

**Table 1:**
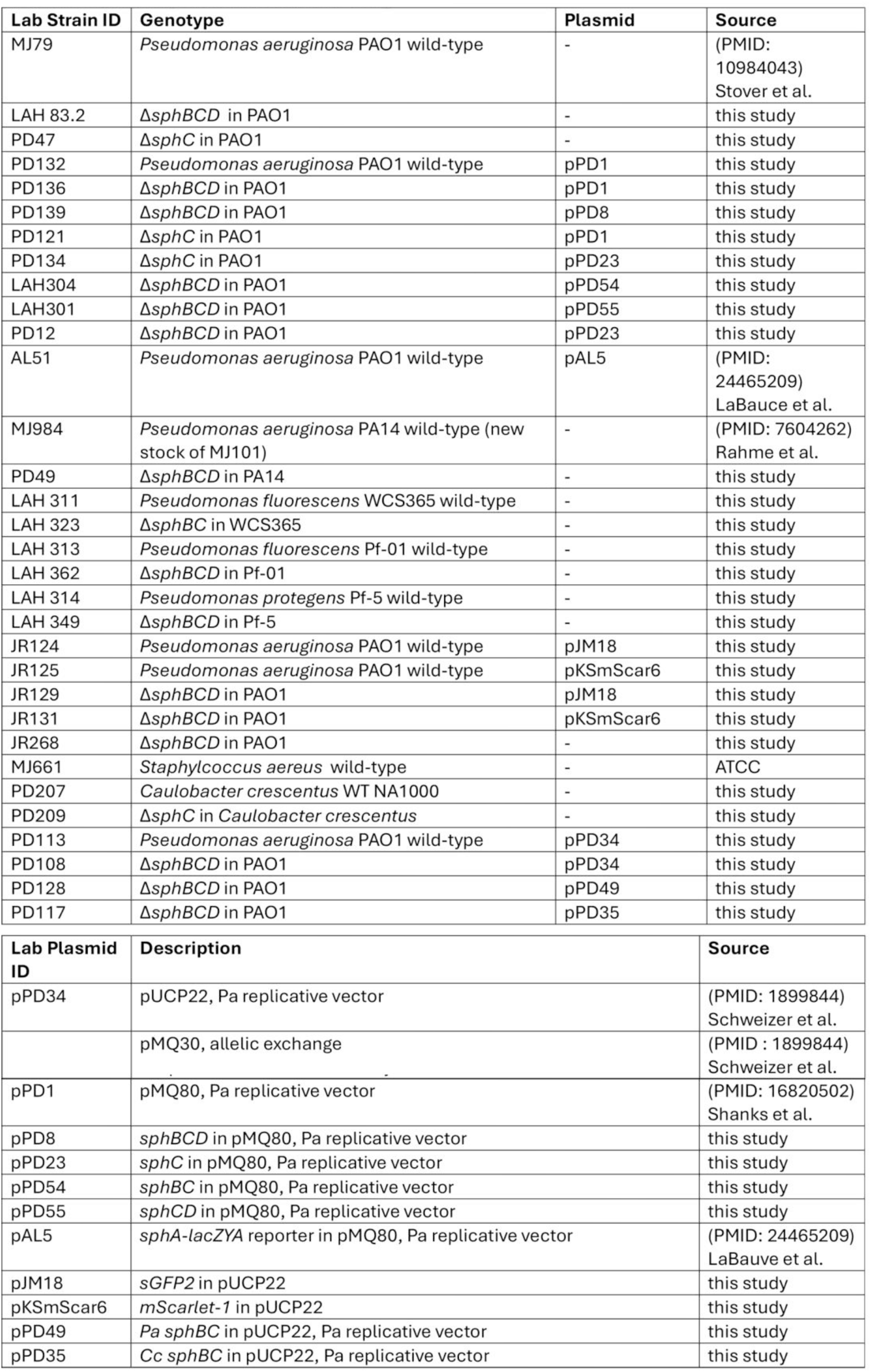
Strains and plasmids used in this study.

| Lab Strain ID | Genotype | Plasmid | Source |
| --- | --- | --- | --- |
| MJ79 | *Pseudomonas aeruginosa* PAO1 wild-type | - | (PMID: 10984043)<br>Stover et al. |
| LAH 83.2 | $\Delta sphBCD$ in PAO1 | - | this study |
| PD47 | $\Delta sphC$ in PAO1 | - | this study |
| PD132 | *Pseudomonas aeruginosa* PAO1 wild-type | pPD1 | this study |
| PD136 | $\Delta sphBCD$ in PAO1 | pPD1 | this study |
| PD139 | $\Delta sphBCD$ in PAO1 | pPD8 | this study |
| PD121 | $\Delta sphC$ in PAO1 | pPD1 | this study |
| PD134 | $\Delta sphC$ in PAO1 | pPD23 | this study |
| LAH304 | $\Delta sphBCD$ in PAO1 | pPD54 | this study |
| LAH301 | $\Delta sphBCD$ in PAO1 | pPD55 | this study |
| PD12 | $\Delta sphBCD$ in PAO1 | pPD23 | this study |
| AL51 | *Pseudomonas aeruginosa* PAO1 wild-type | pAL5 | (PMID: 24465209)<br>LaBauce et al. |
| MJ984 | *Pseudomonas aeruginosa* PA14 wild-type (new stock of MJ101) | - | (PMID: 7604262)<br>Rahme et al. |
| PD49 | $\Delta sphBCD$ in PA14 | - | this study |
| LAH 311 | *Pseudomonas fluorescens* WCS365 wild-type | - | this study |
| LAH 323 | $\Delta sphBC$ in WCS365 | - | this study |
| LAH 313 | *Pseudomonas fluorescens* Pf-01 wild-type | - | this study |
| LAH 362 | $\Delta sphBCD$ in Pf-01 | - | this study |
| LAH 314 | *Pseudomonas protegens* Pf-5 wild-type | - | this study |
| LAH 349 | $\Delta sphBCD$ in Pf-5 | - | this study |
| JR124 | *Pseudomonas aeruginosa* PAO1 wild-type | pJM18 | this study |
| JR125 | *Pseudomonas aeruginosa* PAO1 wild-type | pKSmScar6 | this study |
| JR129 | $\Delta sphBCD$ in PAO1 | pJM18 | this study |
| JR131 | $\Delta sphBCD$ in PAO1 | pKSmScar6 | this study |
| JR268 | $\Delta sphBCD$ in PAO1 | - | this study |
| MJ661 | *Staphylococcus aureus* wild-type | - | ATCC |
| PD207 | *Caulobacter crescentus* WT NA1000 | - | this study |
| PD209 | $\Delta sphC$ in *Caulobacter crescentus* | - | this study |
| PD113 | *Pseudomonas aeruginosa* PAO1 wild-type | pPD34 | this study |
| PD108 | $\Delta sphBCD$ in PAO1 | pPD34 | this study |
| PD128 | $\Delta sphBCD$ in PAO1 | pPD49 | this study |
| PD117 | $\Delta sphBCD$ in PAO1 | pPD35 | this study |
| Lab Plasmid ID | Description | Source |  |
| pPD34 | pUCP22, Pa replicative vector | (PMID: 1899844)<br>Schweizer et al. |  |
|  | pMQ30, allelic exchange | (PMID : 1899844)<br>Schweizer et al. |  |
| pPD1 | pMQ80, Pa replicative vector | (PMID: 16820502)<br>Shanks et al. |  |
| pPD8 | $sphBCD$ in pMQ80, Pa replicative vector | this study | |
| pPD23 | $sphC$ in pMQ80, Pa replicative vector | this study | |
| pPD54 | $sphBC$ in pMQ80, Pa replicative vector | this study | |
| pPD55 | $sphCD$ in pMQ80, Pa replicative vector | this study | |
| pAL5 | $sphA-lacZYA$ reporter in pMQ80, Pa replicative vector | (PMID: 24465209)<br>LaBauve et al. | |
| pJM18 | *sGFP2* in pUCP22 | this study |  |
| pKSmScar6 | *mScarlet-1* in pUCP22 | this study |  |
| pPD49 | *Pa sphBC* in pUCP22, Pa replicative vector | this study |  |
| pPD35 | *Cc sphBC* in pUCP22, Pa replicative vector | this study |  |

### General Allelic Exchange, Chromosomal Alterations, and Electroshock Transformations

All allelic exchanges were completed using the pMQ30 non-replicative and counter-selectable vector(49). Briefly, once constructs were cloned into the pMQ30 backbone, they were transformed into chemically competent S17 λ*pir E. coli* by heat shock. For conjugation, donor and recipient cells were mixed, collected by centrifugation, and resuspended in a small volume of LB and spotted onto LB plates to dry after which they were incubated overnight at 30°C. Single crossover merodiploids were selected by plating on PIA with 50μg/ml gentamicin at 37°C, which also kills the donor *E. coli*. Two independent single crossovers for each allele were inoculated into LB and incubated at 37°C for 3-4 hours with shaking before plating on LB and LB with no NaCl and including 5% sucrose and incubated at 30°C overnight. Sucrose-resistant colonies were then patched to LB with 5% sucrose and no NaCl plates (incubated at 30°C) and LB with 50μg/ml gentamicin plates (incubated at 37°C) to identify and discard remaining merodiploids. Verification of strains from double crossovers was completed using PCR as described.

Allelic exchange vectors for deletion of *sphBCD* or *sphC* in PAO1 and PA14 were created by splice overlap extension as we have described previously for other sphingosine related genes(23). Briefly, PCR fragments were amplified for both upstream and downstream of *sphBCD* or *sphC* and ligated into pMQ30 cut with either KpnI/HindIII or BamHI/HindIII. For *sphBCD* PCR fragment amplification, the upstream region was amplified with primers 2080 and 2083, while the downstream region was amplified using 2081 and 2083. For *sphC* PCR fragment amplification, the upstream region was amplified with primers 1022 and 1024, while the downstream region was amplified using 1023 and 1025. After verification by digest screening, plasmids were sequenced by Plasmidsaurus. Sequence verified plasmids were transformed into chemically competent S17 λ*pir E. coli* and allelic exchange was completed as described above, resulting in strains LAH 83.2 (PAO1 Δ*sphBCD*), PD49 (PAO1 Δ*sphBCD*), and PD47 (PAO1 Δ*sphC*).

Allelic exchange vectors for *sphBCD* deletion in *P. fluorescens* Pf-01 and *P. protegens* Pf-5 and *sphBC* deletion in *P. fluorescens* WCS365 were created using splice overlap extension (SOE) as described above. After amplification and splice overlap, fragments were ligated into pMQ30 cut with KpnI/HindIII (for *P. fluorescens* WCS365 and *P. protegens* Pf-5) or XbaI/KpnI (for *P. fluorescens* PF-01). The *P. fluorescens* WSC365 *sphBC* upstream region was amplified with primers 2736 and 2737, while the downstream region was amplified using primers 2738 and 2739. The *P. fluorescens* Pf-01 *sphBCD* upstream region was amplified with primers 2740 and 2741, while the downstream region was amplified using 2742 and 2743. The P. *protegens* Pf-5 *sphBCD* upstream region was amplified using primers 2732 and 2733, while the downstream region was amplified using 2734 and 2735. After verification by digest screening, plasmids were sequenced by Plasmidsaurus. Sequence verified plasmids were transformed into chemically competent S17 λ*pir E. coli* and allelic exchange was completed as described above, resulting in strains LAH 323 (WSC365 Δ *sphBC*), LAH 362 (Pf-01 Δ*sphBCD*), and LAH 349 (Pf-5 Δ*sphBCD*).

The *sphBCD*, *sphBC*, *sphCD*, and *sphC* complementation constructs, pPD8, pPD54, pPD55, and pPD23, were generated by amplifying the appropriate region from genomic DNA using primer pairs 2726 & 2727, 2882 & 2883, 2726 & 2727, and 2511 & 2512, all cut with EcoRI and HindIII and independently ligated into similarly cut pMQ80. Plasmids with correct insert determined by PCR were sequenced (Plasmidsaurus) and correct plasmids electrotransformed into target strains (**Table 1**). The empty vector control for all complementations was the empty pMQ80 vector.

The *sGFP2* construct, pJM18, and *mScarlet-1* construct, pKSmScar6, were built using HiFi (NEB) assembly using synthetic gene fragments (gBlocks) and ligated into pUCP22 digested with BamHI and EcoRI. pJM18 and pKSScar6 assemblies were verified by digest screening using HindIII and SacI and digest-correct clones were sequenced (Plasmidsaurus) before electrotransformation into target strains (*Table 1*).

### Chemicals and notes on sphingolipid stability, solubility, and handling

All media, media components, and standard chemicals were purchased from ThermoFisher or Sigma. The sphingoid bases sphingosine, phytosphingosine, and sphinganine were purchased from Cayman Chemicals and dissolved in 95% ethanol as aliquots of 50 mM stocks and stored at -20 °C. Storing as aliquots is important, as multiple freeze-thaw cycles lead to loss of each of the sphingoid bases’ antimicrobial capacity and ability to stimulate gene induction via SphR(23). Sphingoid bases were delivered to the vulture vessel in ethanol and then the solvent evaporated to dryness, using air drying for multi-well plastic plates and a gentle stream of nitrogen gas for glass tubes.

### Determining IC_50_ for sphingosine, sphinganine, and phytosphingosine in glass and plastic

*P. aeruginosa* strains were grown overnight at 37 °C shaking in MOPS media 25 mM sodium pyruvate, 5 mM glucose, and 20 μg/ml gentamicin. Cells from overnight cultures were collected via centrifugation, washed with MOPS media, and resuspended in MOPS with 25 mM sodium pyruvate and 20 μg/ml gentamicin. Starting at an OD_600_ of 0.05, Pa strains were grown for 18 hours at 37°C with horizontal shaking in either plastic 48-well plates or 13x100mm glass tubes in the presence or absence of various concentrations of each sphingoid base. For the incubation periods, plates were covered with a sterile, breathable, adhesive microporous sealing film (USA Scientific) to allow for equal gas exchange for each well, while glass tubes were covered loosely with aluminum foil. After 18-hour incubations, OD_600_ was measured using a Synergy H1 (Biotek) plate reader. IC_50_ values calculated in GraphPad Prism using the log(inhibitor) vs response – Variable slope (four parameter) curve fitting analysis.

### Kinetic growth assays

To measure growth kinetics, sphingoid bases were used at 200 μM. Prior to inoculation, *P. aeruginosa* strains were grown overnight at 37 °C, shaking in MOPS media with 25 mM sodium pyruvate, 5 mM glucose, and 20 μg/ml gentamicin. Cells were collected via centrifugation, washed in MOPS media, and resuspended in MOPS with 25 mM pyruvate and 20 μg/ml gentamicin, at a starting OD_600_ of 0.05 in 48-well plates sealed with breathable adhesive films. Absorbance for the film was removed by determining the difference between the absorbance post film application to the read pre-application and subtracting that difference for each well and applying that to all reads for that well. Growth was measured via OD_600_ taken every 30 minutes with a Synergy 2 H1 Biotek hybrid plate reader set at 37°C with orbital shaking before each read.

### Growth assays for other Pseudomonads, *Caulobacter,* and heterologous complementation

To investigate the importance of *sphBCD* in other *Pseudomonas* strains and species, overnight cultures in MOPS media with 25 mM sodium pyruvate and 5mM glucose were grown at 37 °C for *P. aeruginosa* strains and 30 °C for *P. protegens* and *P. fluorescens* strains. Cells were collected via centrifugation, washed in MOPS media, and resuspended in MOPS media with 25 mM pyruvate (or 20 mM pyruvate, 10 mM glucose, or 10 mM succinate when assessing catabolite repression, shown in supplemental figures). *Pseudomonas* strains and species were grown in sterile 13x100mm boroscilicate glass tubes or plastic 48-well plates for 18 hours at 37 °C (for *P. aeruginosa* strains) or 30 °C (for *P. protegens* and *P. fluorescens* strains), with orbital shaking, in the presence or absence of sphingoid bases (200 µM final concentration) at a starting OD_600_ of 0.05. After 18-hour incubations, growth was measured by OD_600_ using a Synergy H1 Biotek plate reader.

*Caulobacter crescentus* WT NA1000 and related Δ*sphC* were maintained at 30 °C on PYE (peptone-yeast extract) plates containing 2 g/L Bacto Peptone, 1 g/L Yeast Extract, 1 mM MgSO_4_, and 0.5 mM CaCl_2_. Prior to assay set up, strains were grown shaking at 30 °C overnight in M2 minimal salts medium (6.1 mM Na_2_HPO_4_, 3.9 mM KH_2_PO_4_, 9.3 mM NH_4_Cl, 0.5 mM MgSO_4_, 10 uM FeSO_4_ (EDTA chelate), and 0.5 mM CaCl_2_) with 0.2% glucose as the sole carbon source. To investigate the importance of *sphBC* in other gram-negative bacterium, such as *C. crescentus*, overnight cultures in M2 minimal media were grown at 30 °C. Cells were collected via centrifugation and resuspended again in M2 minimal media at an OD_600_ of 0.05 and grown in sterile 13 X100 mm borosilicate glass tubes at 30 °C, with orbital shaking, in the presence or absence of sphingosine, at varying concentrations. After 18-hour incubations, growth was measure by OD_600_ using a Synergy H1 Biotek plate reader.

To investigate the importance of homologous *sphBC* from *C. crescentus* in rescuing *P. aeruginosa* Δ*sphBCD* growth inhibition in the presence of sphingosine, overnight *P. aeruginosa* cultures in MOPS media with 25 mM sodium pyruvate, 5 mM glucose, and 20 μg/ml gentamicin were grown shaking overnight at 37 °C. *sphBCD* complementation was assessed with native *P. aeruginosa* genes (PD128; PAO1 Δ*sphBCD* with Pa*sphBCD* on pUCP22) or *C. crescentus* homologues (PD117; PAO1 Δ*sphBCD* with Cc*sphBC* on pUCP22). Cells were collected via centrifugation, washed in MOPS media, and resuspended in MOPS media with 25 mM pyruvate with 20 μg/ml gentamicin. *P. aeruginosa* strains were grown in sterile 13x100mm borosilicate glass tubes for 18 hours at 37 °C, with orbital shaking, in the presence or absence of sphingosine, at a starting OD_600_ of 0.05. After 18-hour incubations, growth was measured by OD_600_ using a Synergy H1 Biotek plate reader.

### *sphA-lacZ* reporter assay

To determine the amount of sphingoid base remaining in culture when *sphBCD* is deleted, *sphA* transcriptional induction was measured using our previously described *sphA-lacZ* reporter assay and construct(23). *P. aeruginosa* was electrotransformed with the *sphA* promoter construct (pAL4)(23), and resultant colonies were grown overnight at 37 °C, shaking, in MOPS media with 25 mM sodium pyruvate, 5 mM glucose, and 20 μg/ml gentamicin prior to induction. Cells were collected by centrifugation, washed in MOPS media, and resuspended in MOPS media with 25 mM sodium pyruvate and 20 μg/ml gentamicin with or without lipid extracts from strains to be tested. Lipid extracts were collected for each strain after 18 hours incubation in the presence or absence of sphingoid bases (200 μM final concentration). β-galactosidase assays were then completed as previously described(50, 51) using Miller’s method(52).

### Thin layer chromatography

To visualize the amount of sphingosine remaining in culture in the presence or absence of *sphBCD,* we used thin layer chromatography. Pa strains were grown overnight at 37 °C, shaking in MOPS media with 25 mM sodium pyruvate, 5 mM glucose, and 20 μg/ml gentamicin. Cells were collected by centrifugation, washed in MOPS media, and resuspended in MOPS media with 25 mM pyruvate and 20 μg/ml gentamicin at a starting OD_600_ of 0.05. Strains were grown for 18 hours at 37 °C, with orbital shaking, in sterile foil-covered borosolicate 13x100mm glass tubes with or without 200 μM sphingosine. After incubation period, lipids were extracted from whole cell culture using the Bligh and Dyer method (53). Briefly, chloroform:methanol (1:2; v:v) was added, samples were vortexed, and one volume of water was added. After briefly vortexing, samples were centrifuged for 10 minutes at 14,000 x g. After centrifugation, the lower organic fraction was collected and dried using N_2_ gas before final resuspension in 20 μL of ethanol. TLC silica gel 60 F_254_ plates (Sigma Aldrich) were pre-run with acetone, dried, and lipid extracts spotted onto the plate. After samples dried, plates were run in a closed glass chamber with chloroform:methanol:water (65:25:4; v:v:v) as the mobile phase. After the mobile phase approached top of the plate, the plate was removed, dried, and was sprayed with Ninhydrin Solution (Acros Organics) to detect sphingosine by its primary amine group.

### LC/ESI-MS/MS

To directly quantify the levels of sphingosine remaining in culture in the presence and absence of *sphBCD*, LC/ESI-MS/MS was completed by Lipotype, Inc (Germany). Strains were grown as per TLC and, after incubation, samples were lysed at 4 °C for 10 minutes via bead beating with vortex cell disruptor using 0.5 mm glass beads. Samples were stored at -80 °C until shipment to Lipotype, Inc. Before LC/ESI-MS/MS, samples were spiked with deuterated internal standards (including 0.25 ng sphingosine-d7). Methanol/isopropanol was added for protein precipitation and the cleared solutions were analyzed using an Agilent 1290 HPLC system with binary pump, multisampler, and column thermostat with a Kinetex EVO C-18, 2.1 x 100 mm, 2.6 μm column using a gradient solvent system of ammonium carbonate (2 mM) and methanol. The flow rate was set at 0.4 mL/min and the injection volume was 1 uL. The HPLC was coupled with an Agilent 6495 Triplequad mass spectrophotometer (Agilent Technologies, Santa Clara, USA) with electrospray ionization source. Analysis was performed with Multiple Reaction Monitoring in positive mode, with at least two mass transitions for each compound. All sphingolipids were calibrated using individual standards. The Agilent Mass Hunter Quant software was used for quantification.

### *P. aeruginosa* Competition Assays

*P. aeruginosa* strains were grown overnight at 37 °C, shaking in MOPS media with 20 mM sodium pyruvate, 5 mM glucose, and 20 μg/ml gentamicin prior to competition assay set up. Cells were collected via centrifugation, washed three times in MOPS media, and resuspended in MOPS media with 20 mM pyruvate and 20 μg/ml gentamicin and normalized to an OD_600_ of 0.5. Sterile borosilicate 13x100 mm glass tubes had vehicle or sphingosine, for a final concentration of 200 µM, dried as described above. To these tubes, 900 mL of MOPS media with 20 mM sodium pyruvate and 20 μg/ml gentamicin was added, followed by 50 µl each of 0.5 OD_600_ GFP and mScarlet expressing *P. aeruginosa* for total starting OD_600_ of 0.05. All cultures were grown at 37 °C, shaking for 18 hours. At 0 and 18-hour timepoints, OD_600_ and GFP (485/528 nm) and mScarlet (550/610 nm) fluorescent signals were measured using a Synergy H1 plate reader (BioTek). Background fluorescence for GFP and mScarlet was corrected by subtracting the signal from WT monoculture carrying the opposite fluorescent protein (mScarlet or GFP, respectively). Corrected fluorescence values were expressed as a percentage of monoculture of WT carrying GFP or mScarlet (set to 100%). Additionally, 20 µL aliquots of each culture were serially diluted in R2B and spot plated onto MOPS media agar plates with 20 mM sodium pyruvate and 5 mM glucose for colony forming unit (CFU) counts at each timepoint. Total colony forming units were counted, GFP-expressing colonies were detected by UV transillumination and appropriate excitation filter and imaged using a ChemiDoc XRS+ Gel Imaging System (BIO RAD). mScarlet expressing colonies were calculated by subtracting GFP-expressing colonies from the total CFU/mL.

### *P. aeruginosa* – *S. aureus* Competition Assays

*P. aeruginosa* was grown overnight in MOPS media with 20 mM sodium pyruvate and 5 mM glucose, shaking at 37 °C. Cells were collected by centrifugation, washed three times with MOPS media, and added to 1 mL MOPS media with 20 mM pyruvate, 5 mM glucose, and 20 µM sphingosine at a final OD_600_ of 0.05 to allow time for *sphBCD* induction. During this incubation, overnight 37 °C LB cultures of *S. aureus* were collected via centrifugation, washed three times with R2B, and adjusted to an OD_600_ of 0.5 in R2B. *P. aeruginosa* diluted into R2B +/- 100 µM sphingosine for one hour, shaking at 37 °C. After one hour, *S. aureus* was added to an OD600 of 0.05 to the P. aeruginosa-containing media or R2B +/- 100 µM sphingosine, and grown for five hours shaking at 37 °C. At 0 and 5 hours of co-culture, 20 µL aliquots serially diluted in R2B, and spot plated onto both PIA and tryptic soy agar (TSA) +7.5% NaCl to select for growth of *P. aeruginosa* and *S. aureus*, respectively, and colony forming unit (CFU) counted.

---

**Supplemental Figure S1:**
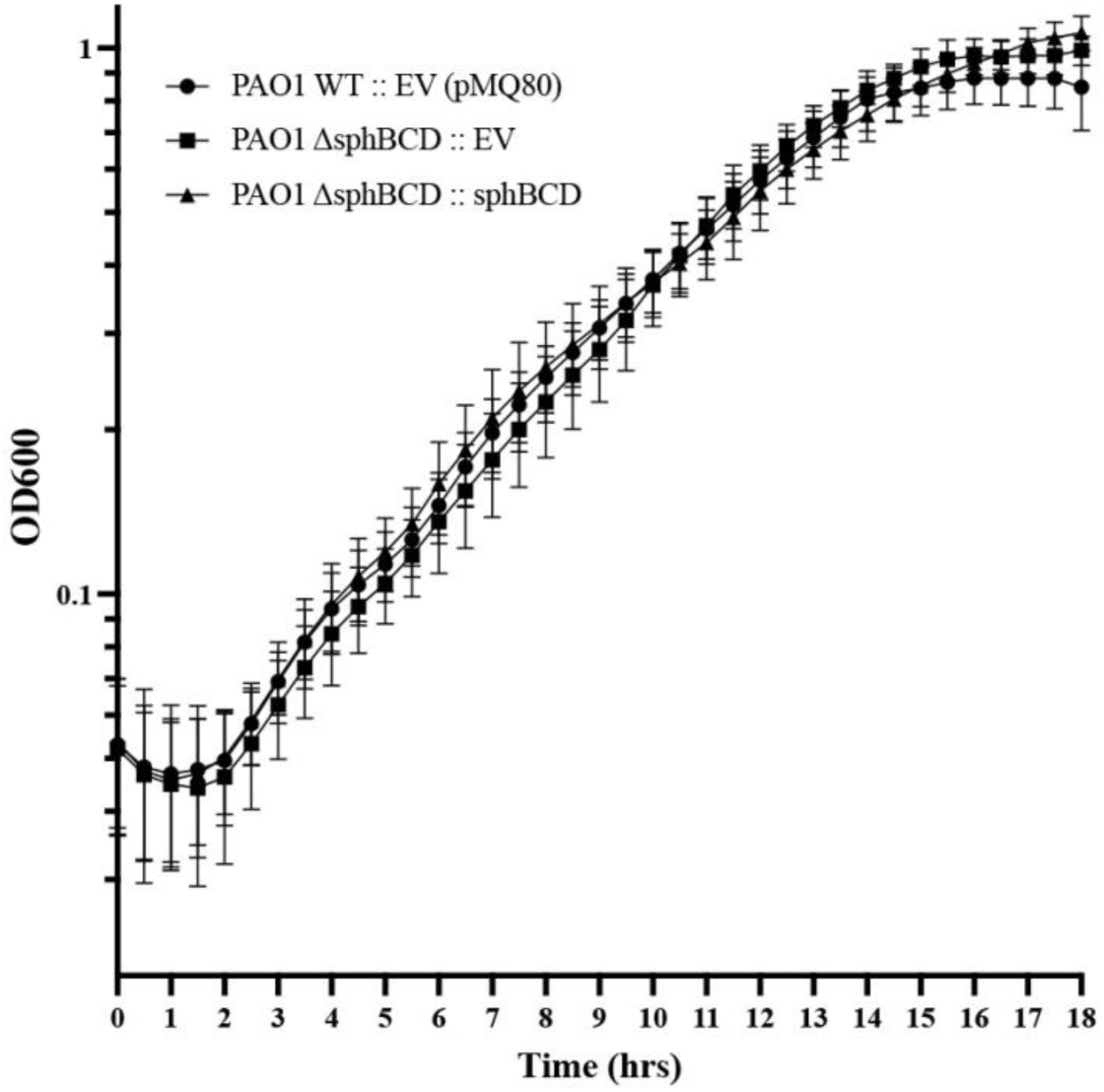
Kinetic growth assessment wild-type, mutant, and complemented strains in the absence of sphingoid bases. Panels shows 18-hour growth timecourse in MOPS pyruvate media measuring growth of each strain by OD_600_. Abbreviations: EV, empty vector pMQ80; sphBCD, vector containing *sphBCD*.

**Supplemental Figure S2:**
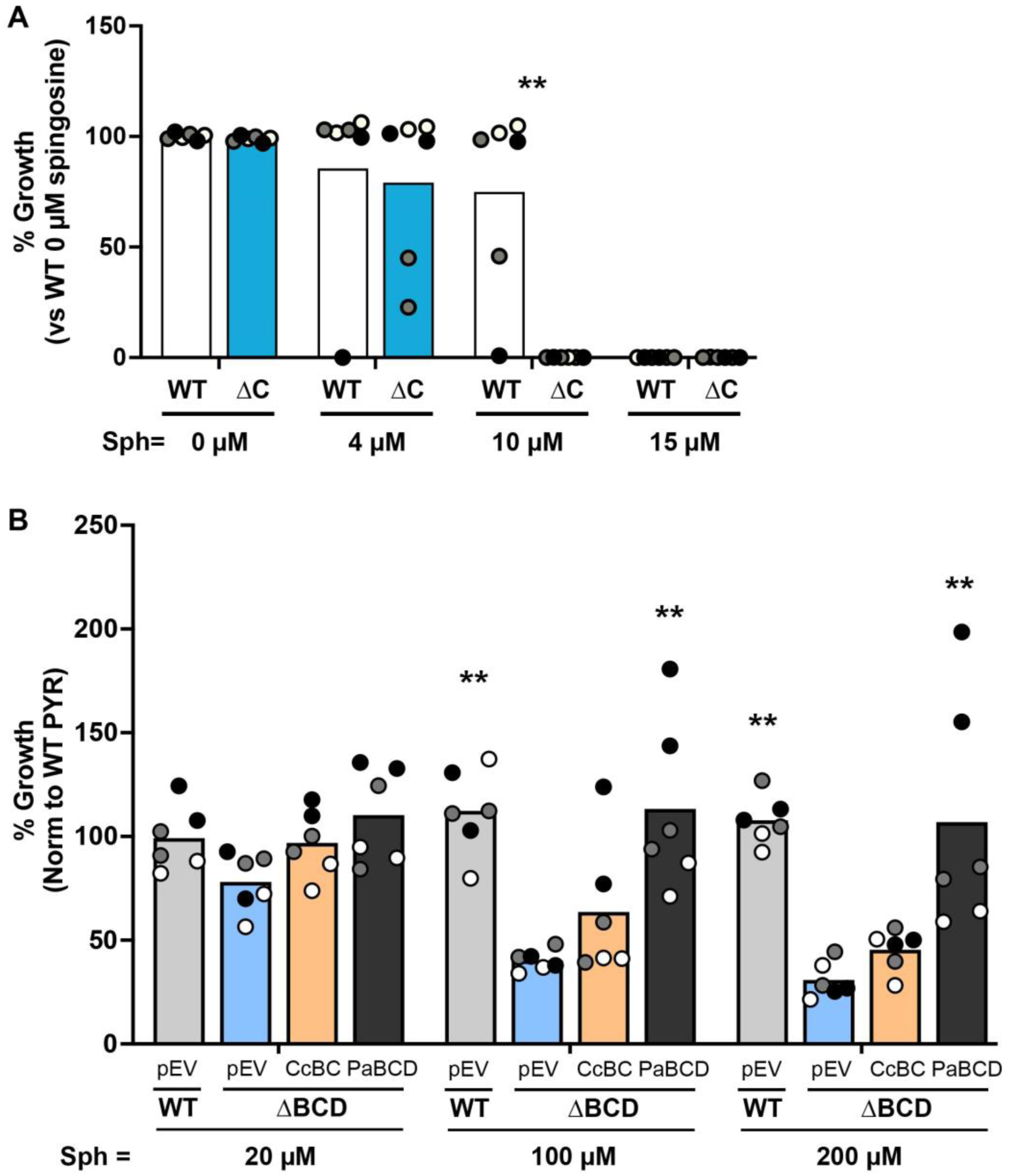
Tests of *Caulobacter crescentus sphBC* function. **(A)** Deletion of the *Caulobacter crescentus sphC* has very little impact on growth in the presence of sphingosine after 18-hour growth in glass when considering the small concentration range of sphingosine over which the effect is noted. **(B)** Complementation analyses of *P. aeruginosa* Δ*sphBCD* with empty vector (pEV), a plasmid containing *C. crescentus sphBC* (CcBC), or a plasmid containing *P. aeruginosa sphBCD* (PaBCD) shows that *C. crescentus sphBC* fails to significantly complement the *P. aeruginosa* Δ*sphBCD* mutant. All data points are shown and are colored by experiment with white circles for replicates from experiment #1, gray from experiment #2, and black from experiment #3. Only the means for each experiment are used in the statistical analyses for these panels (n = 3 per condition). In **A**, significance noted as (**, p<0.01) calculated using multiple Mann-Whitney tests comparing WT to Δ*sphC* within each sphingosine concentration. This test was chosen since the zero growth as a mean within an experiment make the data non-parametric. In **B**, significance noted as (**, p<0.01) calculated from Two-way ANOVA with Dunnett’s post-test comparing each group to the Δ*sphBCD* + pEV group within each concentration. Abbreviations: Sph, sphingosine; Pyr, pyruvate.

**Supplemental Figure S3:**
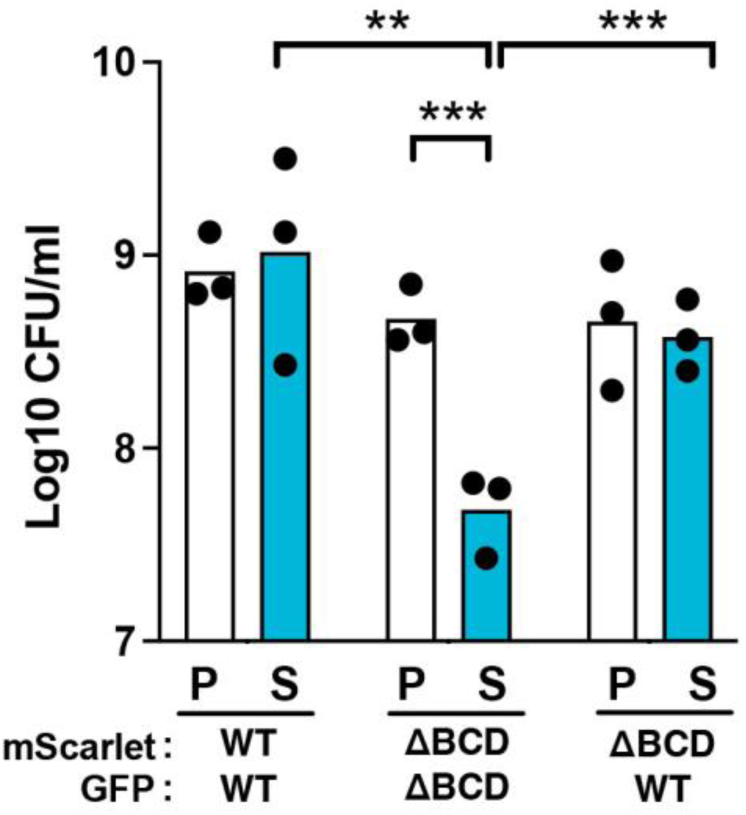
Swap of the fluorescent marker as used in Figure 7. CFU counts of mScarlet-expressing colonies in the presence (S) or absence (P) of sphingosine. The strain carrying each fluorescent protein is labeled below graph. Significance noted as (**, p<0.01; ***, p<0.001) calculated from ANOVA with Tukey’s post-test comparing within and between co-culture groups. Each point represents the mean from a single experiment (n = 3 per condition). Abbreviations: ΔBCD, Δ*sphBCD*; P, pyruvate (control); S, sphingosine; N.D., not detectable.

**Supplemental Figure S4:**
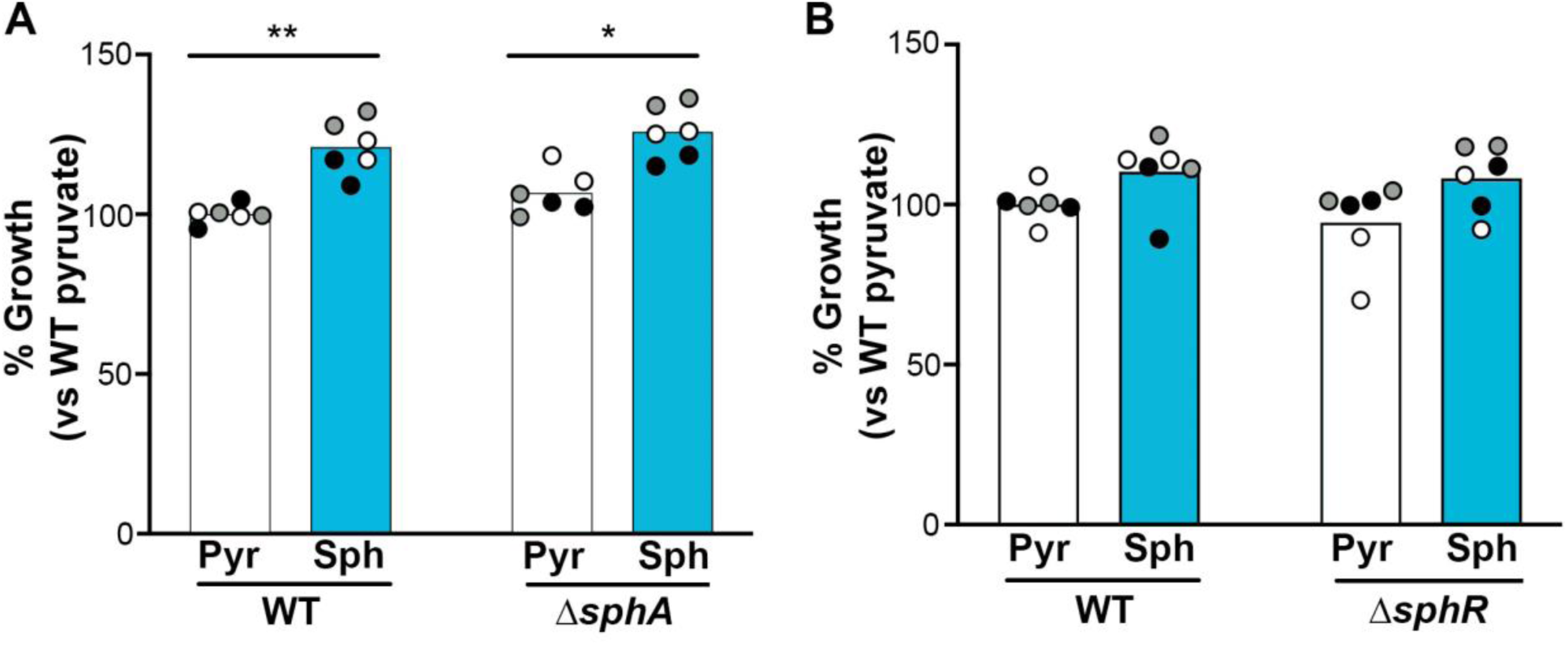
Effects of *sphA* and *sphR* deletion on sphingosine susceptibility. 18-hour growth was normalized to WT growth in MOPS pyruvate set as 100%. **(A)** PAO1 WT compared to the *sphA* deletion strain. **(B)** PAO1 WT compared to the *sphR* deletion strain. For each panel, all data points are shown and are colored by experiment with white circles for replicates from experiment #1, gray from experiment #2, and black from experiment #3. Only the means for each experiment are used in the statistical analyses for these panels (n = 3 per condition). Significance noted as (*, p<0.05; **, p<0.01) calculated from Two-way ANOVA with Sidak’s post-test comparing pyruvate to sphingosine within each strain. Abbreviations: Pyr, pyruvate; Sph, sphingosine.

**Supplemental Figure S5:**
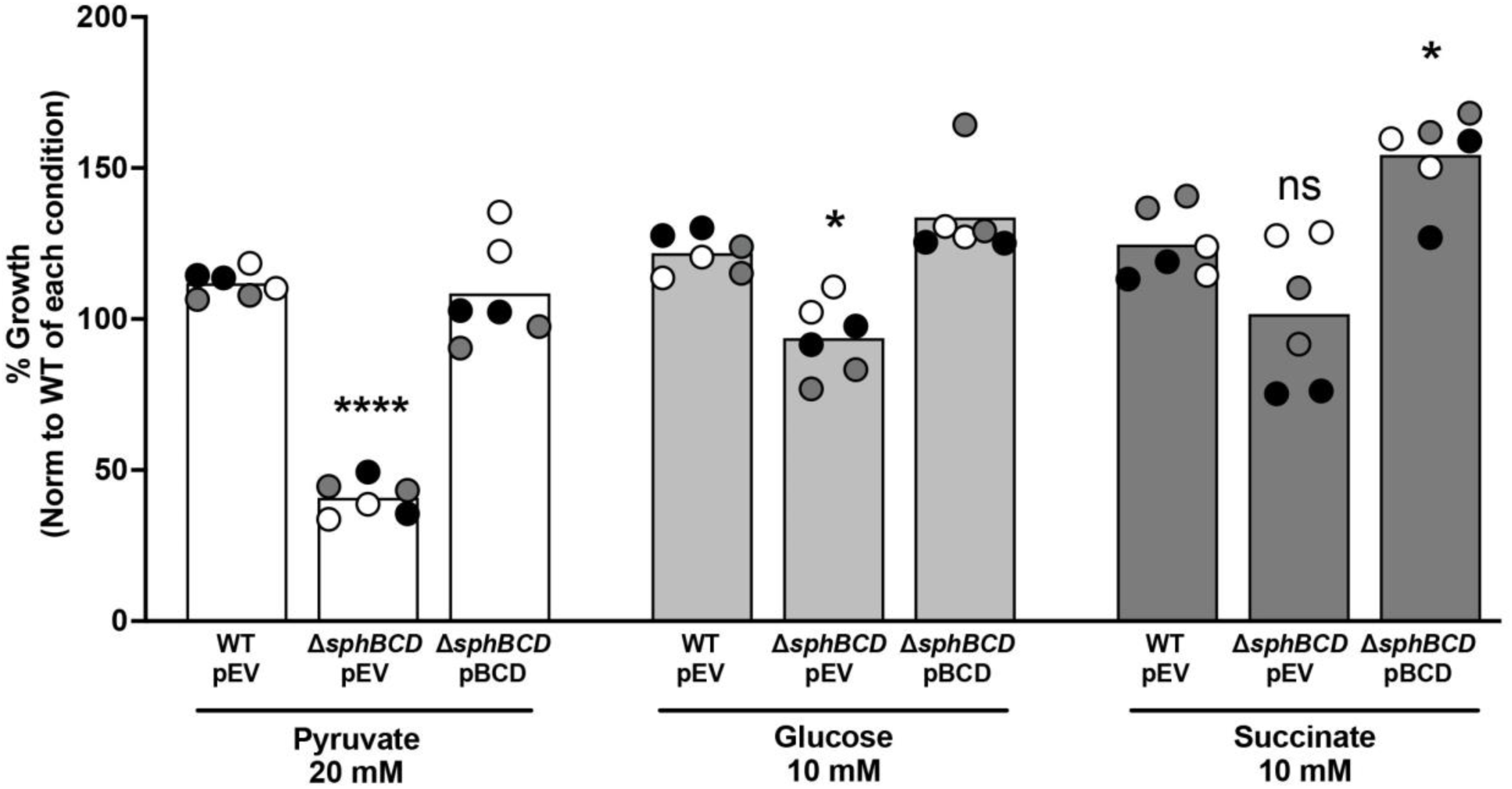
Effect of carbon source on sphingosine inhibition and the role of *sphBCD*. 18-hour growth was normalized to WT+pEV growth in MOPS pyruvate set as 100%. All data points are shown and are colored by experiment with white circles for replicates from experiment #1, gray from experiment #2, and black from experiment #3. Only the means for each experiment are used in the statistical analyses for these panels (n = 3 per condition). Significance noted as (*, p<0.05; ****, p<0.0001) calculated from Two-way ANOVA with Dunnett’s post-test comparing deletion or complement within each carbon source condition to its WT+pEV. Abbreviations: pEV, empty vector pMQ80; pBCD.

**Supplemental Table S1:**
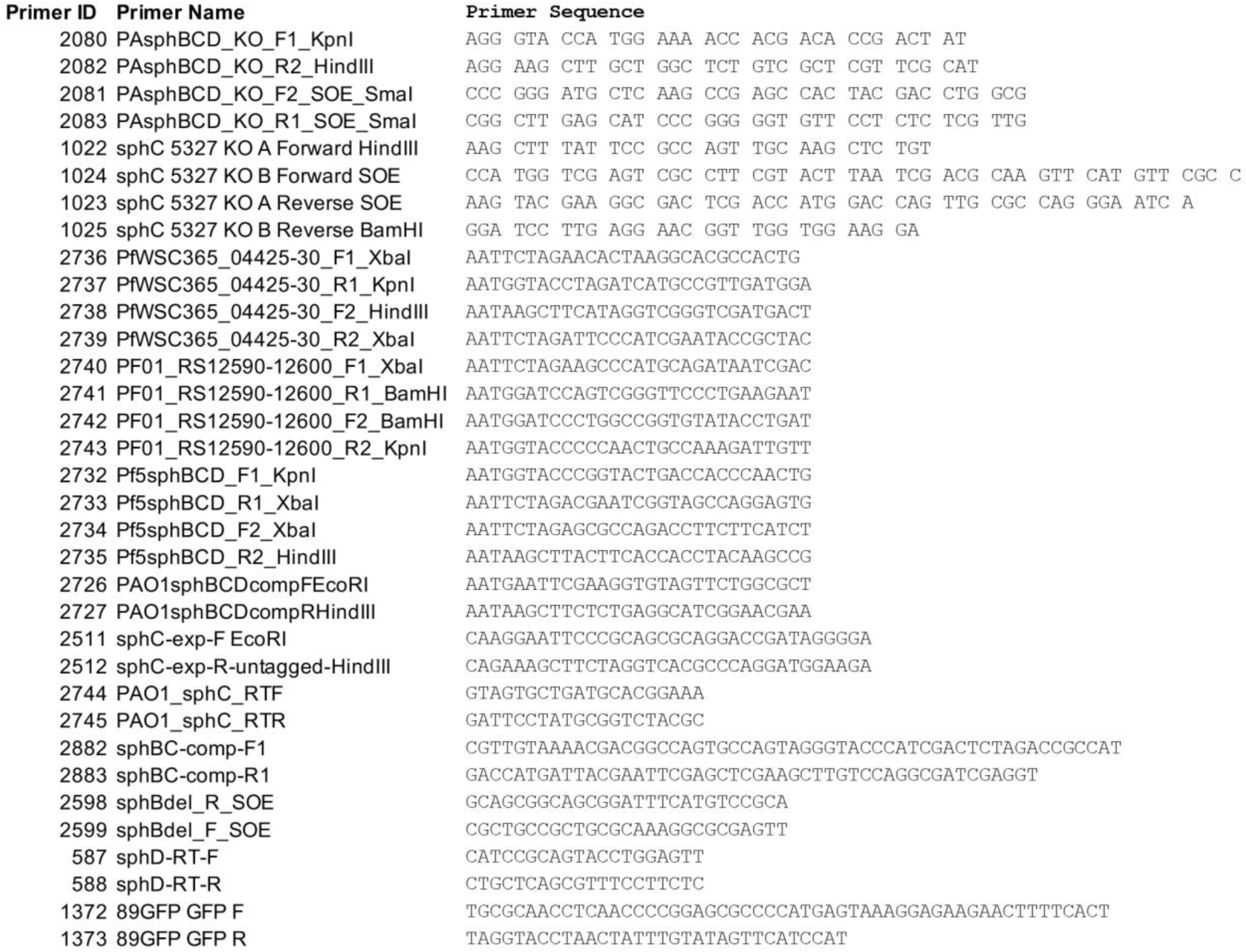
Primers used in this Study.

